# A Novel Human Distal Tubuloid-on-a-Chip Model for Investigating Sodium and Water Transport Mechanisms

**DOI:** 10.1101/2025.03.27.644139

**Authors:** Murillo D L Bernardi, Emre Dilmen, Dorota Kurek, Henriette L. Lanz, Jos Joore, Joost G. Hoenderop, Maarten B. Rookmaaker, Marianne C. Verhaar

## Abstract

The distal segments of the nephron play a central role in regulating water and electrolyte balance, making them critical targets for therapeutic interventions. Damage to these segments is associated with significant health consequences. Studying their (patho)physiology in vivo remains challenging due to the kidney’s complex architecture. Recent advances led to the development of more representative *in vitro* models, including human tubuloids that replicate the phenotype of distal tubule segments. Additionally, novel high throughput microfluidic systems, which support 3D cell culture under flow conditions, provide a platform for closely mimicking *in vivo* environments. This study presents an enhanced in vitro model of human distal tubule segments by integrating tubuloid culture with the OrganoPlate® platform. Tubuloid cells were grown as three-dimensional tubules against a collagen-1 matrix and under alternating flow condition, then differentiated into a distal phenotype. qPCR analysis demonstrated enhanced expression of distal segment markers in 3D flow cultures compared to traditional 2D models. Immunohistochemistry confirmed the formation of a leak-tight, highly polarized epithelium with apical and basolateral localization of key electrolyte transporters. Functional integrity was verified by restricted dextran diffusion and increased transepithelial resistance. Radiolabeled sodium assays revealed active and selective sodium transport mediated by apical epithelial sodium channels (ENaC) and basolateral Na/K ATPase. Sodium transport was followed by water movement, evidenced by dome formation beneath the epithelial monolayer. The model’s utility was further demonstrated in toxicity studies using trimethoprim, an antibiotic that inhibits ENaC function, resulting in reduced sodium transport and dome formation. This system enables the study of primary human tubule cells with a distal phenotype under controlled flow conditions, allowing direct assessment of water and salt transport. The model provides a valuable tool for investigating distal nephron (patho)physiology and facilitates high-throughput drug development and toxicity testing.

## Introduction

The kidney plays a crucial role in maintaining homeostasis through various physiological processes, particularly the regulation of water and salt balance. Within the kidney’s functional units (nephrons), the distal segments are responsible for fine-tuning this regulation^1^. Dysfunction in these segments can lead to severe disturbances in water and salt homeostasis^2–4^. Conversely, these segments serve as important therapeutic targets, such as for diuretic medications that modulate water and salt balance^5^.

Despite their physiological significance, there is a notable lack of suitable models for studying distal segment function. The kidney’s complex architecture makes it challenging to isolate and study specific segments in vivo. Current in vitro models also have significant limitations. Immortalized cell lines, while commonly used, often fail to accurately represent human physiology due to immortalization artifacts and species differences. Additionally, traditional two-dimensional cultures lack crucial physiological features such as apical flow, lack of three dimensional architecture, and interaction with the extracellular matrix, which can significantly influence renal epithelial differentiation and function^6–9^. Furthermore, functional assessments frequently rely on indirect measures of electrolyte transport instead of actual compound transport^10–13^.

Recent advances in tubuloid culture have enabled more representative studies of human distal tubule segments in vitro. Tubuloids are primary human kidney cell cultures derived from urine or biopsy samples that can extensively expand and differentiate into various nephron segments without genetic manipulation^14,15^. We recently published a novel differentiation protocol that enables tubuloid differentiation into phenotypes of the distal nephron^16^. These differentiated tubuloids form leak-tight epithelia in 2D cultures and demonstrate active sodium transport, as verified through radiolabeled sodium uptake studies.

Concurrent developments in high-throughput 3D culture systems have led to significant technological advances, notably the OrganoPlate system from Mimetas. This platform enables the culture of three-dimensional tubules with the capacity for up to 64 parallel experiments per plate and includes flow exposure capabilities^17–19^. The system allows both apical and basal access for measurements and facilitates in situ immunohistochemistry. Our previous work with proximally differentiated tubuloids demonstrated the OrganoPlate’s utility for investigating epithelial barrier integrity and transport mechanisms, including organic cation transporter and P-glycoprotein function^20^. However, these studies were conducted in a lower throughput plate, with proximal phenotype tubuloids and did not examine water and electrolyte transport.

In this study, we aim to establish and characterize distally differentiated tubuloids in the OrganoPlate, evaluating their differentiation through qPCR and immunohistochemistry. We will assess functionality by measuring barrier integrity and directly analyzing electrolyte and water transport mechanisms. Additionally, we will validate the model’s applicability for drug toxicity studies using trimethoprim, an antibiotic known to inhibit sodium uptake in the cortical collecting duct. This work seeks to develop a robust platform for investigating human distal nephron segment (patho)physiology under controlled conditions while enabling high-throughput drug development applications.

## Methods

### Tubuloid culture

Human kidney tubuloid cultures were derived from healthy adult donor tissues from leftover biopsy materials. Tubuloids were isolated from patient kidney biopsies according to the protocol defined by Schutgens *et. al*^15^ and aliquots of passage number 2 were frozen for future use. At the time of the experiment, tubuloids were thawed and seeded in droplets of Cultrex Reduced Growth Factor Basement Membrane Extract (BME, R&D Systems, 3533-010-02) and cultured in medium (ADMEM/F12 supplemented with 1% (v/v) penicillin/streptomycin, HEPES (Thermo Fisher Scientific, 15630056), GlutaMAX (Thermo Fisher Scientific, 35050038), N-acetylcysteine (1⍰mM, Sigma, A9165) and 1.5%(w/v) B27 supplement (50×; serum-free; Thermo Fisher Scientific, 17504044), supplemented with 1%(w/v) Recombinant RSPO3-Fc Fusion Protein conditioned medium (U-Protein Express, R001), Animal-free recombinant human EGF (50⍰ng⍰ml–1, Peprotech, AF-100-15), Recombinant human FGF-10 (100⍰ng⍰ml^−1^, Preprotech, 100-26), Rho-kinase inhibitor Y-27632 (10⍰µM, Abmole, M1817), A8301 (5⍰µM, Tocris Bioscience, 2939) and Primocine (0.1⍰mg⍰ml–1, Invitrogen, ant-pm-1). Tubuloids were cultured at 37 °C, 5%(v/v) CO^2^, medium was refreshed every 2–3 days and they were passaged or harvested 5-7 days after splitting. Cells were kept in culture until passage 10.

Mycoplasma contamination was tested for each donor-derived tubuloid batch following isolation using a colorimetric detection kit (MycoAlert®, Lonza), and only mycoplasma-negative cultures were used for experiments. Sterility was assessed every 48 hours by monitoring culture media for bacteriostasis and fungistasis through visual inspection for turbidity and contamination, ensuring consistent experimental conditions.

### Single cell dissociation and seeding

To obtain a single-cell suspension, tubuloids were treated with dispase II (1 mg/ml, Thermo Fisher Scientific, 17105041) in the tubuloid medium. BME droplets containing tubuloids were resuspended using a 200-μl pipette. The gel-containing medium was then incubated at 37 °C for 30 minutes to allow gel dissolution. The tubuloids were transferred to a 15-ml conical tube, and ice-cold basic medium was added up to 10 ml. Following centrifugation (300g, 4 °C, 5 minutes) and removal of the supernatant, the tubuloid pellets were resuspended in 3 ml Accutase (Thermo Fisher Scientific, 00-4555-56) supplemented with 10 μM Y-27632 dihydrochloride. The tubuloids were sheared approximately 10 times using a p200 tip connected to a 5 mL pipette and using the pipette boy. The suspension was incubated at 37 °C, and every 5 minutes, another 3 mL of warm Accutase was added and shearing was performed again until single-cell dissociation was achieved (about 15 minutes, or 3 times the process). Once single-cell suspension was confirmed, the cells were centrifuged, the supernatant was removed, and the tubuloid-derived cells were resuspended in 1 ml of expansion medium.

### OrganoPlate 3-lane seeding

Tubules were cultured in an OrganoPlate 3-lane 64 based on protocols previously described^20^. Collagen-seeded plates were obtained directly from Mimetas (OrganoReady Collagen 3-lane 64 # MI-OR-CO-CU-02, Mimetas, NL). For cell seeding, the gel inlet wells were cleared of HBSS using the aspiration system and disposable tips. The top medium perfusion channel inlet received 2 µl of the cell suspension (10 × 10^6^ cells/ml) using a repeating single pipette. To ensure comparable cell numbers across chips, the cell suspension was resuspended and a new volume was taken up after every 5-10 chips. Cell-free controls, consisting of 2 µl of cell-free medium, were included in the top medium channel. Expansion medium (50 µl) was directly added to the top medium inlet. The plate was placed on its side in the OrganoPlate stand at an angle of approximately ±75° for 3 hours in a humidified incubator (37°C, 5%(v/v) CO_2_) to allow cell adhesion to the ECM gel. Subsequently, expansion medium (50 µl) was added to the top medium outlet and the bottom medium inlet, followed by addition of expansion medium (50 µl) to the bottom medium outlet. Subsequently, perfusion was initiated by placing the plates in an incubator on an interval rocker (OrganoFlow®, MI-OFPR-L, Mimetas, NL) set at 14° inclination and 8-minute interval. The medium in the perfusion channels was refreshed every 2-3 days to maintain optimal conditions.

Tubuloids were grown in expansion media until the tubules were fully formed (5-7 days), then cultured for 7 days in distal-tubule differentiation media. This differentiation medium, termed FAF medium, consisted of basal media supplemented with three key components: 10 μM forskolin (F6886, Merck) to activate cAMP signalling pathways essential for distal nephron differentiation, 5 μM A83-01 (SML0788, Merck) to inhibit TGF-β signalling and promote epithelial differentiation, and 10 μM fludrocortisone acetate (F0180000, Merck) to activate mineralocorticoid receptor signalling crucial for collecting duct function. This FAF combination specifically drives tubuloid differentiation toward thick ascending limb and collecting duct phenotypes^16^. Tubuloid-derived single cells were cultured in parallel in 24-well plates for comparative gene expression analysis. They were kept in expansion media for 5-7 days, then differentiated in FAF for 7 days.

### Barrier integrity assay (BI assay) and transepithelial electric resistance (TEER)

To evaluate the barrier integrity of tubules grown in an OrganoPlate, we perfused the tubule with a fluorescent dye and observed its radial diffusion across the tubule into the surrounding extracellular matrix (ECM). To perform this assay, a complete cell medium was supplemented with 10 kDa FITC dextran (0.16 mg.mL^−1^, Sigma, DE, FD10S-100MG) and 155 kDa TRITC dextran (0.16 mg.mL^−1^, Sigma, DE, T1287-100MG). The fluorescently labelled medium was then added to the lumen of the tubuloid tubules in the OrganoPlate. The efflux of the dextran molecules from the tubule into the adjacent ECM channel was monitored using a high-content fluorescent microscope (ImageXpress XLS, Molecular Devices, US) equipped with FITC and TRITC channels and a 4× objective.

The barrier integrity was also assessed using a high throughput TEER device (OrganoTEER®, Mimetas, NL). It consists of a plate holder, electrode board with 480 stainless steel electrodes arranged in pairs, a measurement unit with impedance analyzer, and a laptop running dedicated software. The electrode pairs were inserted into the microfluidic channels of the OrganoPlate, forming conductive paths across each chip through the ECM interfaces. The measurement unit acquired impedance spectra from 0.1 Hz to 1 MHz, and the software optimized measurement parameters based on expected TEER values. Single-point TEER measurements involved removing the OrganoPlate from the incubator, refreshing it with 50 μL of cell culture medium, and allowing it to equilibrate at room temperature for 30 minutes before conducting measurements.

### Expression analysis

For expression analysis, total RNA from tubuloids on the OrganoPlate (3D) or cultured as monolayers (2D) was extracted using the RNeasy Mini kit (Qiagen, DE, 74104) following the manufacturer’s protocol. Reverse transcription was then performed using either the SuperScript™ IV First-Strand Synthesis System (Thermo Fisher, US, 18091050) or M-MLV reverse transcriptase (Thermo Fisher, US, 28025013) with random primers (Invitrogen, US, 48190011) according to the respective manufacturer’s protocols. Quantitative PCR was conducted on a Roche Lightcycler 96 using the FastStart Essential DNA Green Master (Roche, CH, 06402712001). Data analysis was carried out using the accompanying software (LightCycler® 96, Roche, CH). All gene expression data were normalized to GAPDH as the housekeeping gene and expressed as fold change relative to the control condition using the 2^(−ΔΔCt) method. Statistical analysis was performed to compare gene expression across different conditions, with results presented as fold changes normalized to control conditions for each donor. Primers used for qPCR are listed below in Table 1.

**Table 1.**
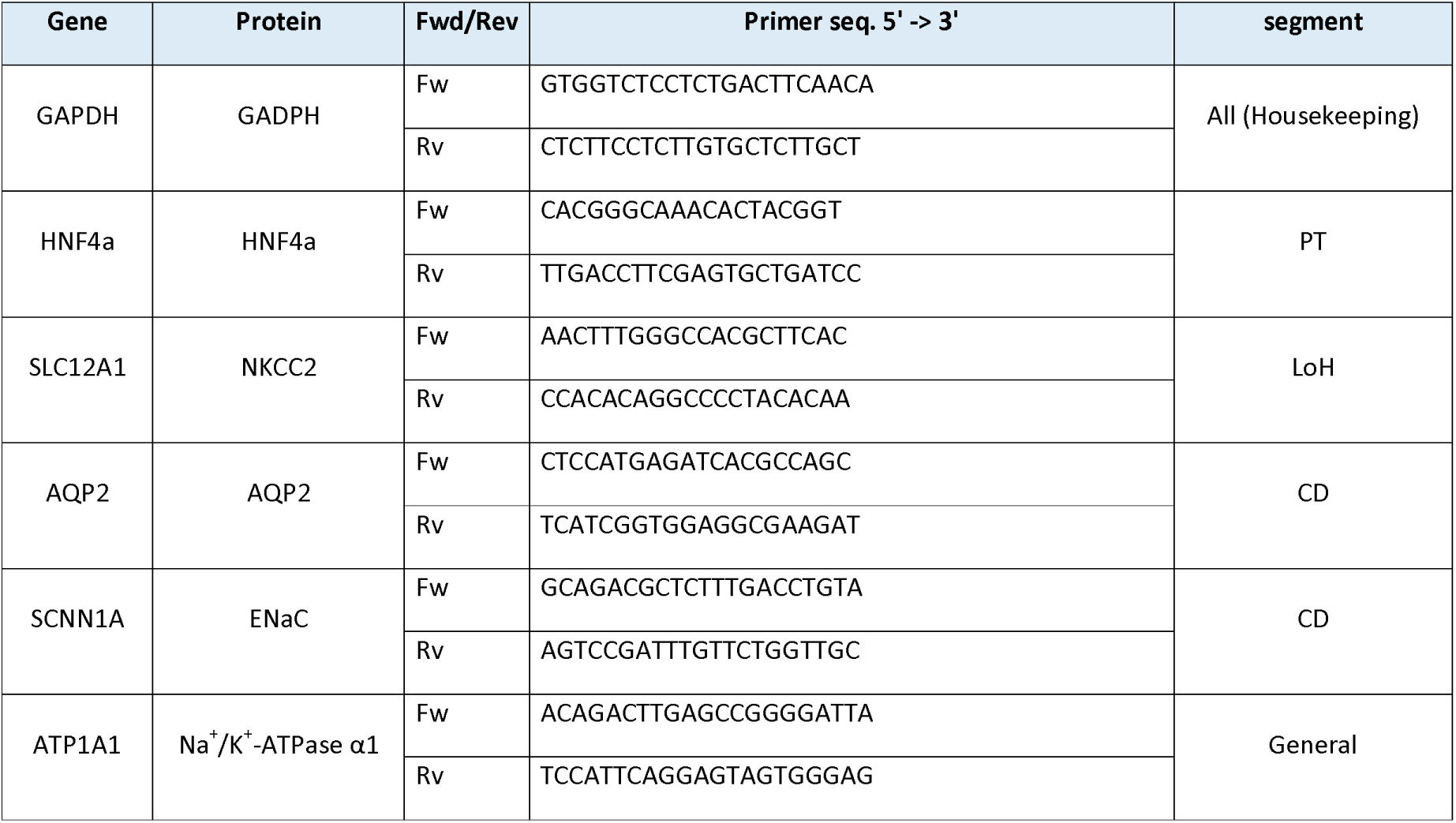
Forward (Fw) and Reverse (Rv) primers for the target genes. PT = Proximal tubule, LoH = Loop of Henle, CD = Collecting duct.

### Immunocytochemistry

Following fixation, immunofluorescent staining was performed on the OrganoPlate cultures. Briefly, cells were permeabilized with Triton X-100 solution and blocked with a buffer containing FBS, bovine serum albumin, and Tween-20. Primary antibodies (**Table 2**) were applied and incubated for 1-2 hours or overnight on a rocking platform, followed by incubation with secondary antibodies for 1 hour. The primary antibodies used are described on the table below. Nuclei were stained using Hoechst dye (ThermoFisher, H3570). Confocal high content imaging microscopy was performed (Micro XLS-C, Molecular Devices) on the stained OrganoPlate using a 10x magnification, capturing images at 3 µm intervals along the microfluidic channel. In addition, laser scanning microscopy (LSM900, Zeiss) with objective 63x (NA 1.4) was used to acquire images and processed with FIJI software (ImageJ).

**Table 2.** Primary and secondary antibodies used in the stainings.

|  | Anti- | Vendor | Source | Dye | Cat nr | Dilution |
| --- | --- | --- | --- | --- | --- | --- |
| **Primary Ab** | Acetylated tubulin | Merck | Mouse | - | T6793 | 1:200 |
|  | E-Cadherin | Cell Signaling Tech | Rabbit | - | 3195S | 1:200 |
|  | ZO-1 | Invitrogen | Rabbit | - | 339100 | 1:50 |
|  | Na<sup>+</sup>K<sup>+</sup>-ATPase | Kind gift from Prof. Koenderink (Radboud University, Nijmegen, the Netherlands)<sup>21</sup> | Rabbit IgG | - |  | 1:200 |
|  | NKCC2 | MRC PPU | Sheep | - | S838B | 1:300 |
|  | AQP2 | Santa Cruz Biotechnology | Rabbit | - | sc-515770 | 1:300 |
| **Secondary Ab** | Rabbit | Life technologies | Goat | Alexa Fluor 488 | RA11008 | 1:250 |
|  | Mouse | Life technologies | Goat | Alexa Fluor 555 | A21422 | 1:250 |
|  | Sheep | Thermo fisher scientific | Donkey | Alexa Fluor 594 | A-11016 | 1:250 |

### Functional sodium uptake assays

The tubuloids were prepared for functional sodium uptake assays by 5-7 days growth until confluency in expansion medium followed by 7 days of either basal medium or differentiation medium. Tubuloids were pre-incubated in hypotonic buffer (70 mM Na^+^-D-gluconate, 2.5 mM K^+^-D-gluconate, 0.5 mM CaCl_2_, 0.5 mM MgCl_2_ and 2.5 mM HEPES, with pH set to 7.4 using Tris) for 30 minutes including inhibitors ouabain (0.1 mM, 102541, MP Biomedicals), hydrochlorothiazide (0.1 mM, H4759, Sigma Aldrich), bumetanide (0.1 mM, B3023, Sigma Aldrich), amiloride (0.1 mM, A7410, Sigma Aldrich) and/or dimethyl sulfoxide (DMSO) as vehicle according to specific conditions. Following the pre-incubation, the tubuloids were exposed to buffers including corresponding inhibitors and radioactive tracer sodium (^22^Na, (U.S. Department of Energy Isotope Program)) for 30 minutes. At the end of the uptake experiment, the tubuloids were washed 3 times with ice-cold hypotonic buffer and consecutively lysed using 10% (w/v) SDS to sample the intracellular contents including the radioactive tracer sodium. The samples were mixed in Opti-Fluor (6013199, Perkin Elmer) fluid and the tracer ^22^Na radioactivity was measured using a liquid scintillation counter (Hidex 300SL).

### Statistical analysis

Statistical analysis was conducted using Prism version 10.4 (GraphPad, United States). For qPCR experiments, samples from 5 chips per donor were pooled to create an individual sample. We compared 3 independent samples for the differentiation condition with their respective expansion condition for all 4 donors. Results are presented as fold-change normalized to control conditions for each donor. For functional assays, 8 individual chips per condition and donor were used, with experiments performed independently for all 4 donors and repeated twice per donor. The data displayed represent the average of these two independent experiments. Comparisons were made within each donor by comparing the experimental condition with its respective negative control. No averaging across donors was performed due to biological variability.

One-sample t-tests were employed to assess deviations from the hypothetical mean of 1 for gene expression comparisons. Additionally, one-way ANOVA combined with Dunnett’s multiple comparisons test or two-way ANOVA combined with Sidak or Tukey multiple comparisons test was used for further statistical analysis. The mean ± standard error of the mean (SEM) is represented by error bars. Statistical significance is indicated by asterisks as follows: *p ≤ 0.05; **p ≤ 0.01; ***p ≤ 0.001; ****p ≤ 0.0001. p values exceeding 5% were deemed not statistically significant (ns) and were excluded from reporting.

### Data Management

Data generated in this study were managed in compliance with the FAIR (Findable, Accessible, Interoperable, and Reusable) principles to ensure transparency and reproducibility. Metadata and experimental details were documented using standard ontologies and formats, facilitating interoperability and reuse. All datasets and analysis scripts have been stored in publicly accessible repositories, with persistent identifiers and detailed documentation provided for easy retrieval. In alignment with the CARE (Collective Benefit, Authority to Control, Responsibility, and Ethics) principles, donor data were handled with strict adherence to ethical guidelines and regulatory standards. Informed consent was obtained, and personal identifiers were removed to ensure confidentiality. Additionally, the use of donor materials was conducted with consideration of collective benefit and respect for the autonomy of donors and their communities.

## RESULTS

### Tubuloids form polarized 3D tubules in the OrganoPlate under flow

To establish a 3D in vitro microfluidic tubuloid model we employed the 3-lane OrganoPlate platform, consisting of 64 independent microfluidic chips, all equipped with three microfluidic channels that converge at the center of each chip, allowing for microscopic monitoring of tubuloid culture (**Figure 1A**). This platform was selected for its ability to provide controlled flow conditions while enabling high-throughput experimentation across multiple biological replicates.

**Figure 1.**
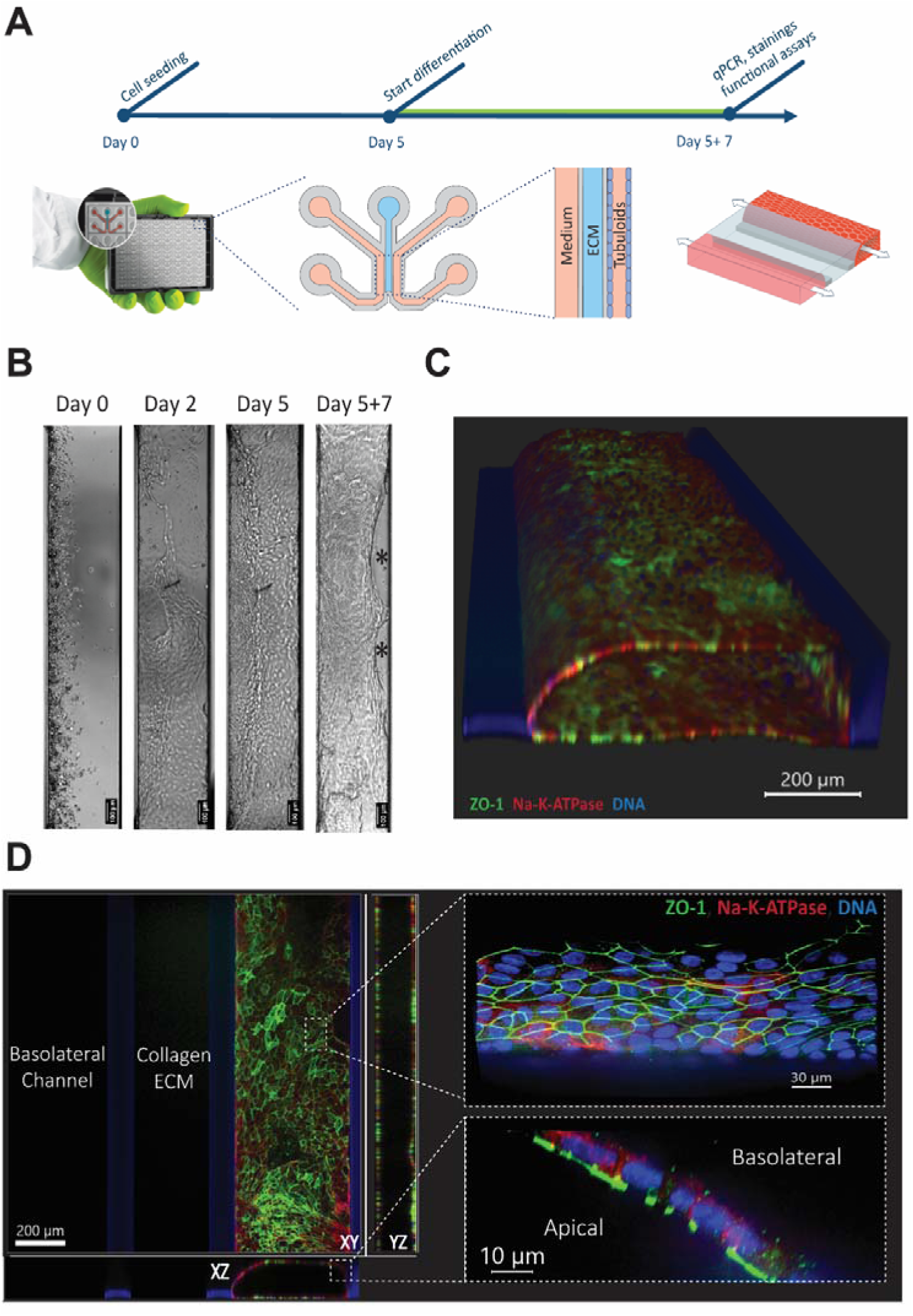
Development and characterization of tubuloid-on-chip model. **(A)** Experimental timeline and platform schematic. Timeline shows cell seeding (Day 0), followed by growth period (Day 5), and subsequent functional analysis (Day 5+7). The OrganoPlate system features three compartments: two perfusable channels flanking a central ECM gel chamber, enabling tubuloid formation and culture. **(B)** Brightfield microscopy showing progressive tubule development: initial cell seeding (Day 0), early attachment (Day 2), and mature tubule formation with visible doming (Day 5+7). Asterisks (*) indicate formation of subepithelial domes (representative images, n=64 chips). **(C)** Immunofluorescence 3D reconstruction demonstrating tubule polarization and barrier formation. **(D)** Confocal projections of a tubuloid chip in the 3-lane microfluidic chip stained for ZO-1 (green), Na^+^/K^+^-ATPase (red), and nuclei (blue). Panels display XY, YZ, and XZ projections, illustrating the epithelial monolayer’s polarization and spatial organization. The right panel shows a magnified view with a cross-section highlighting the polarized epithelial layer.

The seeding approach involved dispersing tubuloid-derived single cells into chips containing patterned extracellular matrix (ECM) gel, creating a structured environment that promotes three-dimensional tubular formation. Within 5 to 7 days of growth under alternating flow conditions, a well-defined tubular structure emerged (**Figure 1B-C**). Following differentiation with FAF medium for 7 days, mature tubules displayed characteristic subepithelial dome formation (**Figure 1B**, asterisks), a phenomenon indicative of active fluid transport across the epithelial barrier, and discussed later in the manuscript. The resulting structure was characterized by a tubule with properly polarized cells, as indicated by basolateral expression of Na+K+-ATPase (**Figure 1D**), demonstrating successful establishment of epithelial polarity essential for physiological transport functions.

### Organoplate flow culture further enhance differentiation towards distal nephron phenotypes

Treatment with FAF medium (forskolin, A 83-01, and fludrocortisone acetate) induced tubuloid differentiation toward thick ascending limb (TAL) and collecting duct (CD) phenotypes, evidenced by increased expression of segment-specific markers SLC12A1, SCNN1a and AQP2 respectively (**Figure 2A**). This differentiation cocktail, which targets cAMP signalling, TGF-β inhibition, and mineralocorticoid receptor activation, effectively promoted the expression of key transporters and channels characteristic of distal nephron segments. Key transporters showed substantial upregulation across donors: NKCC2 (26.9±7.8-fold, *p* < 0.05), ENaC (2.7±1.1-fold, *p* < 0.01), AQP2 (14.2±22-fold *p* < 0.05), and Na+K+-ATPase (2.5±0.5-fold *p* < 0.05). Immunocytochemistry confirmed elevated NKCC2 and AQP2 protein expression in differentiated chips, with distinct population patches (**Figure 2B**). While ENaC upregulation was observed at the mRNA level, specific antibody staining proved challenging.

**Figure 2.**
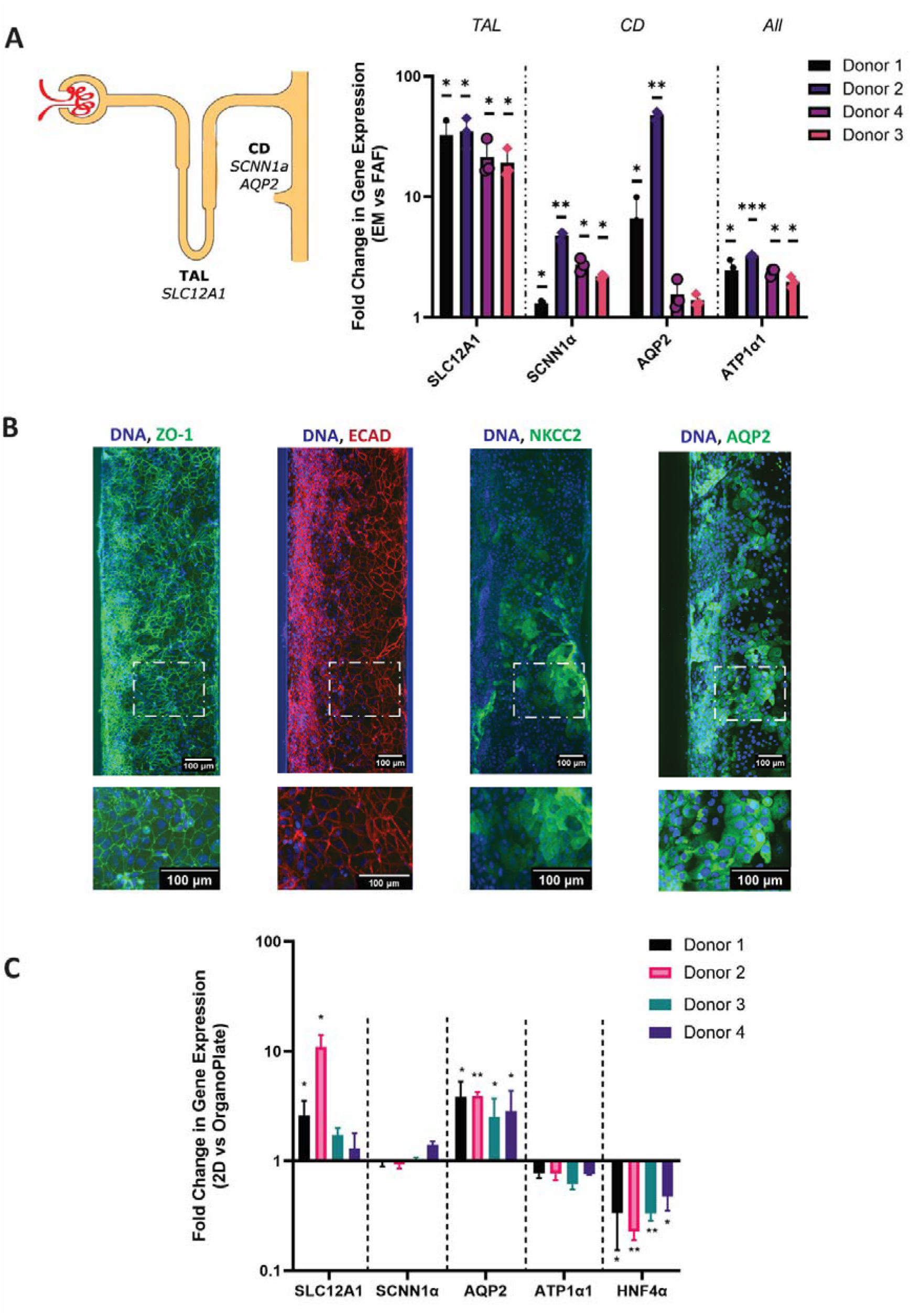
Expression profile of distal nephron markers in differentiated tubuloids. (A) Left: Schematic illustration of distal nephron segments highlighting the thick ascending limb of the loop of Henle (TAL) (SLC12A1/NKCC2) and collecting duct (CD) (SCNN1a/ENaC, AQP2) markers. Right: qPCR analysis showing segment-specific gene expression across four donors after differentiation (fold change relative to basal medium, log scale, Mean ± SD). Statistical analysis was performed using a one-sample t-test comparing fold changes to the hypothetical mean of 1 for each gene. Asterisks indicate statistical significance: *p<0.05, **p<0.01, ***p<0.001. Samples from 5 chips per donor were pooled to create each replicate, with 3 independent samples per condition for each of the 4 donors. EM = Expansion media, FAF = Forskolin, A83-01, Fludrocortisone (distal differentiation media). (B) Immunofluorescence analysis of tubuloid cultures (20X magnification). Upper panels show full-length tubules; lower panels show magnified regions (dotted boxes). Staining shows expression of ZO-1 (tight junctions), ECAD (distal tubule), NKCC2 (TAL), and AQP2 (CD), all counterstained with DNA (representative images, n=2-3). (C) Comparative gene expression analysis between 3D (OrganoPlate) and 2D (tubuloids grown on static 24-well plates) culture conditions across four donors (fold change, n=3). Comparisons made within each donor due to variability in baseline expression. All qPCR data were normalized to the housekeeping gene GAPDH and expressed as fold change relative to the corresponding 2D static culture condition for each donor. Statistical significance was determined using a paired t-test between 3D and 2D conditions, with asterisks indicating significance levels as above.

Differentiation of tubuloids in 3D culture in the microfluidic chip was maintained or even enhanced compared to static 2D culture conditions. To isolate the specific effects of three-dimensional architecture and alternating flow, we compared tubuloids cultured in FAF medium under both conditions. Enhanced differentiation in the 3D platform was observed for NKCC2 (4.1 ± 4.5-fold, *p* < 0.05) and AQP2 (3.3 ± 0.7-fold, *p* < 0.05), whereas expression of ENaC and Na^+^K^+^-ATPase remained comparable to 2D culture expression across donors (**Figure 2C**). Additionally, tubuloids-on-chip showed reduced expression of HNF4α, indicating diminished proximal tubular progenitor characteristics. This reduction in HNF4α expression suggests that the 3D microfluidic environment promotes more complete differentiation away from the progenitor state compared to static 2D culture, supporting enhanced maturation toward the distal nephron phenotype.

### Differentiation increased leak tightness of tubuloids epithelium cultured in the Organoplate

As leak tightness is crucial for renal epithelial cell function in the distal parts of the nephron we analysed leak tightness by histology, diffusion and transepithelial electrical resistance (TEER). Immunostaining revealed ZO-1 expression, indicating tight junction formation (**Figure 2B**). Barrier function was confirmed through dextran-FITC permeability assays (70kD and 155kD), which demonstrated restricted diffusion of molecules in well-formed epithelial monolayers (**Figure 3A**). TEER, the gold standard for barrier assessment, was measured consistently from day 5 through day 5+7 post-differentiation using the OrganoTEER device (**Figure 3B**). FAF differentiation significantly enhanced TEER values after 4 days of differentiation (**Figure 3C).** When evaluating TEER change upon differentiation, we observe an increase in three donors: Donor 1 (54±11% increase, p<0.0001), Donor 2 (88±27% increase, p<0.0001), and Donor 3 (77±44% increase, p<0.0001). Donor 4 showed a modest increase (11±17%, p=0.33) (**Figure 3D**).

**Figure 3.**
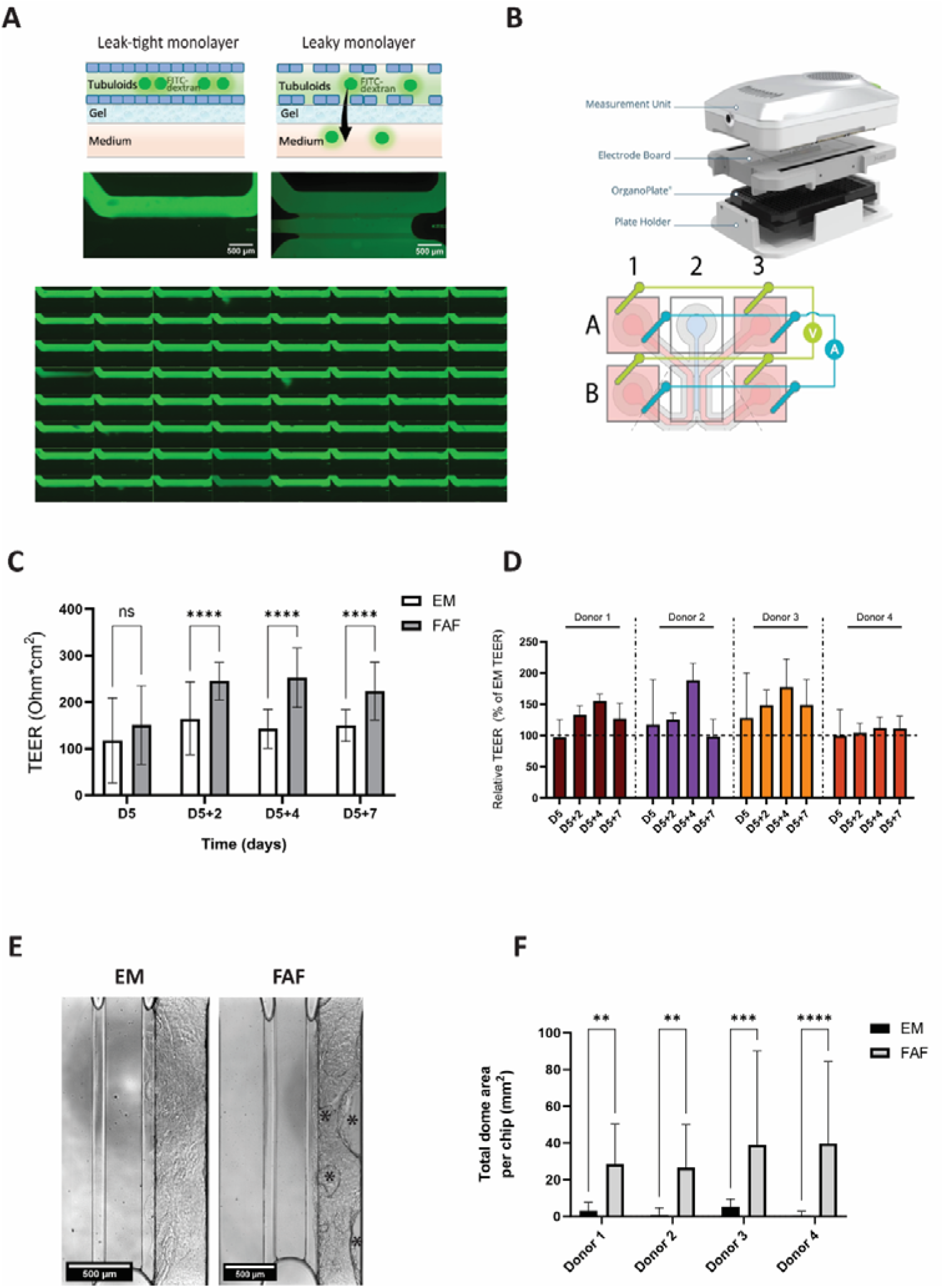
Barrier function and dome formation in tubuloid cultures. **(A)** Dextran-FITC permeability assay demonstrating leak-tight (left) versus leaky (right) tubules, with corresponding schematic representations. Below, a full 3 lane-64 plate is shown on day D5 with all chips composed of leak-tight tubes. **(B)** OrganoTEER measurement system schematic showing device components and electrode configuration. Top: Exploded view of measurement assembly. Bottom: Well arrangement showing current (blue) and voltage (green) electrode pairs for four-point measurements. **(C)** TEER values for donor 1 comparing EM and FAF differentiated chips at different timepoints (n=32, Mean ± SD) **(D)** Time-course of relative TEER measurements across four donors, shown as percentage of expansion media (EM) baseline values (Days 0-7, n=32, Mean ± SD). **(E)** Representative brightfield images comparing dome formation between EM and FAF-differentiated conditions. Asterisks (*) indicates domes in the chip. **(F)** Quantification of total dome area per chip in EM versus FAF conditions (n=16, Mean ± SD, **p<0.01, ***p<0.001, ****p<0.0001).

### Subepithelial dome formation indicates transepithelial water transport by differentiated tubuloids

Differentiated tubuloids exhibited subepithelial dome formation across all donors (**Figure 3E**). Quantitative analysis revealed substantial increases in dome-covered area following FAF differentiation compared to expansion media (EM) across all donors: Donor 1 (3.1±4.5 to 28.5±21.8 mm²), Donor 2 (0.8±3.9 to 26.6±23.5 mm²), Donor 3 (5.2±4.2 to 38.9±51.1 mm²), and Donor 4 (0.6±2.3 to 39.7±44.5 mm²; all n=16, p<0.001) (**Figure 3F**).

### Differentiated tubuloids exhibit transporter-specific sodium transport in the OrganoPlate

To assess sodium reabsorption, a key function of distal nephron segments, we performed radioactive tracer (^22^Na) experiments (**Figure 4A-C)**. Treatment with the basolateral Na+K+-ATPase inhibitor ouabain significantly increased accumulation of intracellular sodium across all donors (p<0.0001, n=8): Donor 1 (27.5±7.1 to 57.4±11.3), Donor 2 (17.0±2.0 to 44.3±9.8), Donor 3 (30.7±5.6 to 81.6±10.3), and Donor 4 (34.2±8.2 to 83.2±28.4 counts per minute) (**Figure 4B)**.

**Figure 4.**
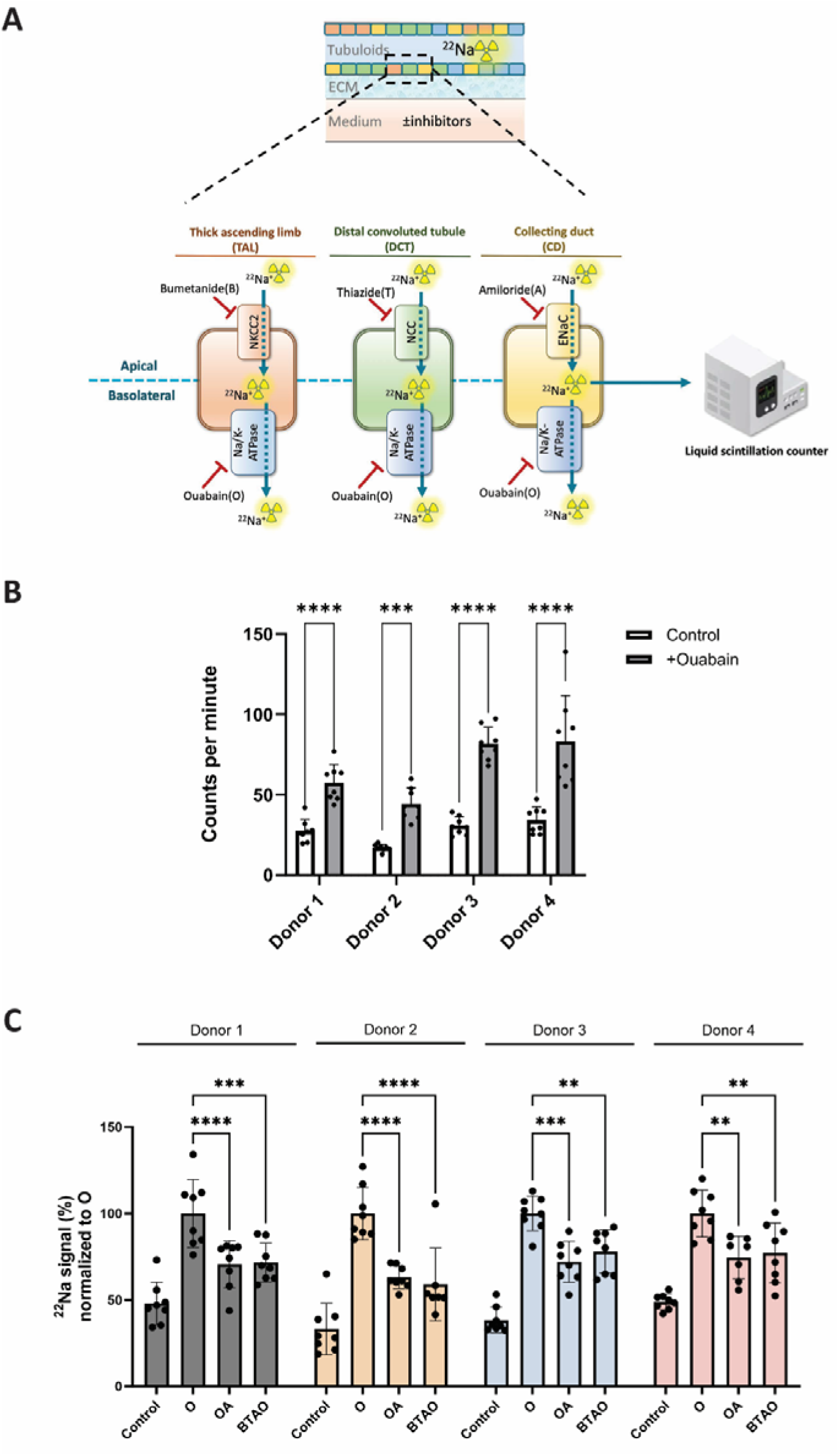
Segment-specific sodium transport analysis in tubuloid cultures. **(A)** Experimental schematic of sodium transport pathway analysis. Diagram shows segment-specific inhibitors targeting distinct transporters: bumetanide (NKCC2 in TAL), thiazide (NCC in DCT), amiloride (ENaC in CD), and ouabain (Na+/K+-ATPase). ^22^Na uptake measured via liquid scintillation counting. **(B)** Baseline versus ouabain-treated intracellular sodium levels across four donors (counts per minute, Mean ± SD) **(C)** Comparative analysis of sodium transport inhibition, normalized to ouabain condition. Treatment groups: ouabain (O), ouabain+amiloride (OA), bumetanide+thiazide+amiloride+ouabain (BTAO). n=8 chips per condition, with experiments performed for all 4 donors and repeated twice, Mean ± SD, **p<0.01, ***p<0.001, ****p<0.0001. Statistical comparisons made within each donor between experimental conditions and respective negative controls.

Subsequently, we sought to pinpoint the specific transporters responsible for this sodium influx by using different inhibitors that block its apical uptake. Bumetanide (B) for NKCC2, thiazide (T) for NCC, and amiloride (A) for ENaC were pre-incubated for 30 minutes on the chips in conjunction with ouabain (O) (**Figure 4A**). Notably, it was only in the presence of ouabain, leading to intracellular sodium accumulation, that we could observe the impact of BTA. As a result, the BTAO condition significantly reduced intracellular sodium accumulation in all donors, reaching up to 40% reduction compared to the O condition (p < 0.01).

To identify the specific transporters involved, we pre-incubated samples for 30 minutes with individual inhibitors followed by 30-minute ouabain exposure. While bumetanide-ouabain (OB) and thiazide-ouabain (OT) combinations showed no significant effect (data not shown), amiloride-ouabain (OA) treatment significantly reduced intracellular sodium accumulation across all donors, comparable to BTAO conditions: Donor 1 (70±13%), Donor 2 (63±6%), Donor 3 (72±11%), and Donor 4 (74±12%) of control values (**Figure 4C**).

### Investigation of Distal Tubular Drug Toxicity beyond viability

We investigated trimethoprim’s (TMP) effects on both cell viability and epithelial sodium channel (ENaC) functionality using our distal-tubuloids-on-chip model. TMP can reduce sodium transport through ENaC inhibition, likely due to its structural similarity to amiloride (**Figure 5A**). To assess potential cytotoxicity, we exposed the tubuloids to high-dose TMP (500 µM) for 72 hours. This prolonged exposure did not affect either transepithelial electrical resistance (TEER) values or cell viability as evaluated by the Alamar Blue assay (**Figure 5B-C**). In contrast, acute TMP exposure significantly reduced ^22^Na accumulation in two of four tested donors: Donor 2 showed a 21% reduction (p<0.05) and Donor 4 exhibited a 27% reduction (p<0.01). These reductions were comparable to amiloride-induced effects (**Figure 5D**).

**Figure 5.**
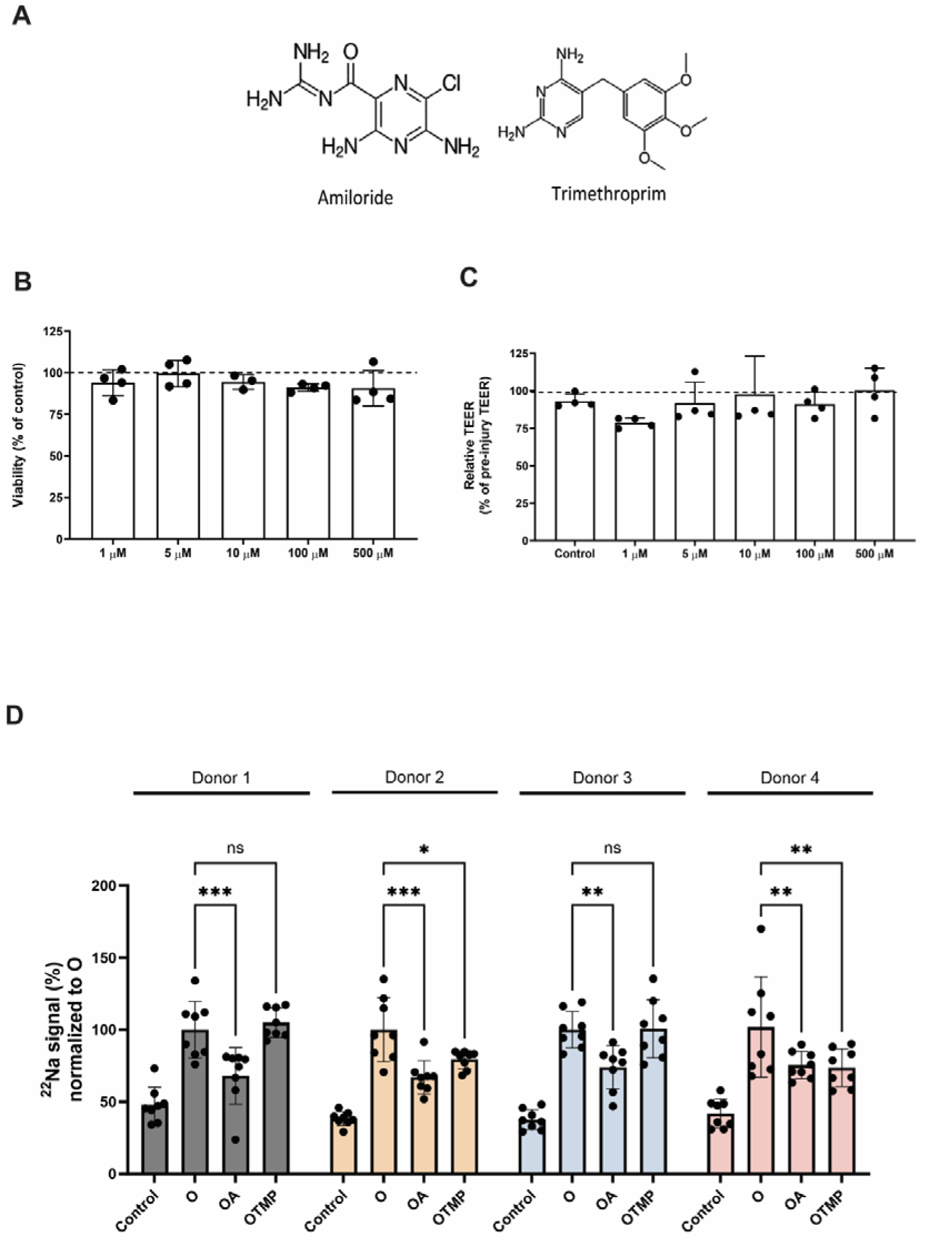
Validation of the distal-tubuloids-on-a-chip model using trimethoprim as a model compound. **(A)** Chemical structures of amiloride and trimethoprim (TMP), highlighting their structural similarity. **(B)** Cell viability assessment of distal tubuloids after 72-hour exposure to increasing TMP concentrations (1-500 µM), showing maintained viability across all concentrations. **(C)** Transepithelial electrical resistance (TEER) measurements following 72-hour TMP exposure, demonstrating preserved barrier integrity. **(D)** ^22^Na uptake measurements in four independent donors comparing the effects of ouabain (O), ouabain+amiloride (OA), and ouabain+TMP (OTMP). TMP significantly reduced sodium uptake in donors 2 and 4, comparable to amiloride’s effect. Data shown as mean ± SD, with 8 individual chips per condition and donor. Experiments performed independently for all 4 donors and repeated twice. Statistical comparisons made within each donor between experimental conditions and respective negative controls. Statistical significance: ns = not significant, *p<0.05, **p<0.01, ***p<0.001.

## DISCUSSION

Our study presents the first successful implementation of human kidney tubuloids with a distal nephron phenotype in a three-dimensional microfluidic platform. Culture in the 3D microfluidic system further enhanced tubuloid differentiation, evidenced by increased expression of functional markers and decreased expression of progenitor cell markers. Immunohistochemistry demonstrated the formation of a polarized leak tight epithelium with the physiological localization of transporters. Functional assessment confirmed leak tightness via diffusion and TEER assays, while demonstrating actual sodium and water transport that could be inhibited by diuretics. Additionally, we successfully replicated drug toxicity effects beyond viability using trimethoprim, showing the reduction of cell function.

Innovative functional 3D kidney models are reshaping our understanding of renal physiology and pathology^22–24^. Most models have focused on the proximal tubule and nephrotoxicity assessment, achieved through methods like hollow fibre systems^25,26^ and 3D printing^27,28^. Meanwhile, some studies have extended the scope, including OrganoPlate-incorporated models using primary kidney cells^17,29–32^ and proximal tubule-differentiated tubuloids^15,20^. In contrast, the functional exploration of human derived-distal nephron segments within 3D models has remained limited.

Our model advances kidney-on-chip technology through several key features. The use of human donor-derived tubuloids enables physiological cell proliferation and expansion, avoiding the metabolic and division alterations observed in immortalized cells^9,33,34^. The introduction of flow and 3D tubular architecture enhanced TAL and CD marker expression compared to 2D cultures. Additionally, the OrganoPlate platform provides accessible apical and basolateral compartments and enables efficient multi-chip experimentation. While our first tubuloid study employed the 3L-40 format^20^, the current model utilizes the 384-well plate design with the 3L-64 format, a more automation-friendly setup that streamlines experimentation and increases compatibility with high-throughput systems. The inclusion of TAL and CD segments, typically underrepresented in kidney-on-a-chip models^24^, combined with demonstrated electrolyte transport functionality through radioactive tracer studies and diuretic responses, establishes this model’s capacity to replicate physiological renal processes.

Recent progress in functional evaluation of distal nephron segments comes from electrophysiological methods applied to 2D models of kidney cells derived from iPSC-organoids. Shi et al. introduced an innovative protocol, guiding iPSCs through stages to generate functional UB organoids and CD cells, indirectly demonstrating amiloride-sensitive ENaC-mediated function in a 2D transwell model through voltage clamping^35^. Montalbetti et al. similarly investigated ion channel functionality within kidney organoid-derived cells, identifying a cell subset with amiloride-sensitive ENaC currents^36^. While these studies provided valuable insights into CD function, they relied on low throughput 2D models and indirect measurements of transport activity. Our model advances beyond these approaches by enabling direct measurement of sodium transport mechanisms in a 3D microfluidic system. Sodium uptake experiments revealed that the effect of the apical transport inhibitor was only detectable in the presence of ouabain, indicating Na^+^K^+^-ATPase’s primary role in trans-epithelial sodium transport regulation over NKCC2 or ENaC. In addition, combined amiloride and ouabain treatment significantly reduced sodium uptake across all donors, suggesting collecting duct segment enrichment. While other transporters were also expressed, only amiloride consistently inhibited ^22^Na uptake, potentially due to variable transporter-expressing cell populations and the distinct kinetics of ENaC as a channel versus NKCC as a transporter^37–39^. This might imply that sodium reabsorption by these transporters is more complex than previously thought, stressing the importance of reductionist models to evaluate actual transport as compared to assessment of indirect parameters like qPCR or protein expression.

Water transport was evaluated in our experiments indirectly by quantifying dome formation. Doming is a prevalent phenomenon observed in kidney epithelial cells cultivated on impermeable substrates, and stems from the movement of water and salt^40–42^. These domes form as fluid is transported beneath the cell monolayer, resulting in the focal detachment of cells from the surface. This phenomenon has been described in distal nephron cells such as Madin-Darby canine kidney (MDCK) cells^43^ and in collecting duct cells such as mpkCCD, where doming was suggested due to ENaC-mediated Na^+^ reabsorption followed by subsequent water absorption^44^. The observation of domes on the OrganoPlate has also been shown in intestinal tubes, where it was linked to active fluid transport across a leak-tight barrier^45^. In our study, we observed the emergence of domes on the plastic side of more than 90% of all differentiated chips. This observation suggests that the upregulation of distal nephron transporters increased electrolyte transport and subsequently led to water transport following from the lumen to the basolateral side of the tubule.

Our model also facilitated an investigation into the potential toxic effects of TMP on electrolyte transport. TMP, a commonly used antibiotic, has been clinically associated with conditions such as hyponatremia and hyperkalemia^46–50^. This effect may be attributed to TMP’s structural similarity to the diuretic amiloride, which can block ENaC channels in the kidney, reducing sodium reabsorption and potassium secretion in the distal nephron ^5 1^. While high-dose treatment did not affect cell viability, it resulted in reduced sodium reabsorption in two of four donors, aligning with clinical observations of impaired tubular function. This donor variability likely reflects natural differences in cell activity among individuals or variations in cell type composition.

While our platform represents a significant advance, several limitations present opportunities for future development. The current heterogeneous population of cell types from different nephron segments, while beneficial for studying cell-cell interactions, can complicate the investigation of cell-type-specific processes. This limitation could be addressed through segment-specific inhibitors or cell sorting techniques. Additionally, the OrganoPlate system’s requirement for bidirectional flow deviates from physiological unidirectional fluid movement, which could be addressed with novel alternative plate designs.

Beyond our current findings, our distal-tubuloids-on-a-chip model offers promising paths for future research. Exploring transport mechanisms of other electrolytes like K^+^, Ca^2+^, and HCO3^−^ within the tubular segments will deepen our insights into renal function. Other commonly used clinical drugs like calcineurin inhibitors may also have adverse effect on electrolyte transport^52^, which could be investigated in such model. Incorporating patient-derived samples can advance personalized medicine, especially for genetic diseases impacting the CD and TAL, such as Liddle and Bartter syndromes. Patient-specific tubuloids can mimic disease conditions, enabling the study of underlying mechanisms and potential treatments. Additionally, integrating endothelial cells into an adjacent compartment could create a dynamic co-culture system, enhancing physiological relevance given the crucial role of endothelial-epithelial communication in kidney function and disease^53–55^.

In conclusion, our 3D kidney tubuloid platform represents a significant advancement in kidney tissue engineering, offering improved physiological relevance and expanded capabilities for drug development and disease modeling. The combination of primary human cells, 3D architecture, and flow conditions creates a more representative model of kidney tubule function, while the observed donor variations provide valuable insights into patient heterogeneity.

## Acknowledgments

The Research Project is financed by the PPP Allowance made available by Top Sector Life Sciences & Health to the Dutch Kidney Foundation to stimulate public-private partnerships within the framework of RegMed XB. The authors also acknowledge the funding from the EU H2020 research and innovation programme under Marie S. Curie Cofund RESCUE grant agreement No 801540.

## References

1. Féraille E, Doucet A. Sodium-potassium-adenosinetriphosphatase-dependent sodium transport in the kidney: hormonal control. Physiol Rev. 2001;81(1):345–418. doi:10.1152/PHYSREV.2001.81.1.345

2. Reilly RF, Ellison DH. Mammalian distal tubule: Physiology, pathophysiology, and molecular anatomy. Physiol Rev. 2000;80(1):277–313. doi:10.1152/PHYSREV.2000.80.1.277/ASSET/IMAGES/LARGE/9J0100060013.JPEG

3. Qian Q, Qi Qian C. Salt, water and nephron: Mechanisms of action and link to hypertension and chronic kidney disease. Nephrology (Carlton). 2018;23(Suppl Suppl 4):44. doi:10.1111/NEP.13465

4. Sebastian A, Hulter HN, Kurtz I, Maher T, Schambelan M. Disorders of distal nephron function. Am J Med. 1982;72(2):289–307. doi:10.1016/0002-9343(82)90822-1

5. Pearce D, Manis AD, Nesterov V, Korbmacher C. Regulation of distal tubule sodium transport: mechanisms and roles in homeostasis and pathophysiology. Pflugers Arch. 2022;474(8). doi:10.1007/s00424-022-02732-5

6. Ko B, Mistry AC, Hanson L, et al. A new model of the distal convoluted tubule. Am J Physiol Renal Physiol. 2012;303(5). doi:10.1152/ajprenal.00139.2012

7. Liu M, Deng M, Luo Q, Dou X, Jia Z. High-Salt Loading Downregulates Nrf2 Expression in a Sodium-Dependent Manner in Renal Collecting Duct Cells. Front Physiol. 2020;10. doi:10.3389/fphys.2019.01565

8. Gartzke D, Fricker G. Establishment of optimized MDCK cell lines for reliable efflux transport studies. J Pharm Sci. 2014;103(4):1298–1304. doi:10.1002/JPS.23901

9. Maqsood MI, Matin MM, Bahrami AR, Ghasroldasht MM. Immortality of cell lines: challenges and advantages of establishment. Cell Biol Int. 2013;37(10):1038–1045. doi:10.1002/CBIN.10137

10. Eladari D, Chambrey R, Peti-Peterdi J. A New Look at Electrolyte Transport in the Distal Tubule. Annu Rev Physiol. 2011;74:325. doi:10.1146/ANNUREV-PHYSIOL-020911-153225

11. Mironova E, Bugay V, Pochynyuk O, Staruschenko A, Stockand JD. Recording Ion Channels in Isolated, Split-Opened Tubules. Methods Mol Biol. 2013;998:341. doi:10.1007/978-1-62703-351-0_27

12. Lourdel S, Paulais M, Marvao P, Nissant A, Teulon J. A Chloride Channel at the Basolateral Membrane of the Distal-convoluted Tubule a Candidate ClC-K Channel. Journal of General Physiology. 2003;121(4):287–300. doi:10.1085/JGP.200208737

13. Reilly RF, Ellison DH. Mammalian distal tubule: Physiology, pathophysiology, and molecular anatomy. Physiol Rev. 2000;80(1):277–313. doi:10.1152/PHYSREV.2000.80.1.277/ASSET/IMAGES/LARGE/9J0100060013.JPEG

14. Yousef Yengej FA, Jansen J, Rookmaaker MB, Verhaar MC, Clevers H. Kidney organoids and tubuloids. Cells. 2020;9(6):1–20. doi:10.3390/cells9061326

15. Schutgens F, Rookmaaker MB, Margaritis T, et al. Tubuloids derived from human adult kidney and urine for personalized disease modeling. Nat Biotechnol. 2019;37(3):303–313. doi:10.1038/s41587-019-0048-8

16. Yousef Yengej FA, Pou Casellas C, Ammerlaan CME, et al. Tubuloid differentiation to model the human distal nephron and collecting duct in health and disease. Cell Rep. 2024;43(1). doi:10.1016/j.celrep.2023.113614

17. Marschner JA, Martin L, Wilken G, Melica ME, Anders HJ. A “Kidney-on-the-Chip” Model Composed of Primary Human Tubular, Endothelial, and White Blood Cells. Methods in Molecular Biology. 2023;2664:107–121. doi:10.1007/978-1-0716-3179-9_8

18. Bonanini F, Kurek D, Previdi S, et al. In vitro grafting of hepatic spheroids and organoids on a microfluidic vascular bed. Angiogenesis. 2022;25(4):455–470. doi:10.1007/S10456-022-09842-9/FIGURES/4

19. Mori A, Vermeer M, van den Broek LJ, et al. High-throughput Bronchus-on-a-Chip system for modeling the human bronchus. Scientific Reports 2024 14:1. 2024;14(1):1–13. doi:10.1038/s41598-024-77665-3

20. Gijzen L, Yousef Yengej FA, Schutgens F, et al. Culture and analysis of kidney tubuloids and perfused tubuloid cells-on-a-chip. Nature Protocols 2021 16:4. 2021;16(4):2023–2050. doi:10.1038/s41596-020-00479-w

21. Koenderink JB, Swarts HGP, Willems PHGM, Krieger E, De Pont JJHHM. A Conformation-specific Interhelical Salt Bridge in the K+ Binding Site of Gastric H,K-ATPase. Journal of Biological Chemistry. 2004;279(16). doi:10.1074/jbc.M400020200

22. Faria J, Ahmed S, Gerritsen KGF, Mihaila SM, Masereeuw R. Kidney-based in vitro models for drug-induced toxicity testing. Arch Toxicol. 2019;93(12):3397–3418. doi:10.1007/s00204-019-02598-0

23. Irvine AR, van Berlo D, Shekhani R, Masereeuw R. A systematic review of in vitro models of drug-induced kidney injury. Curr Opin Toxicol. 2021;27. doi:10.1016/j.cotox.2021.06.001

24. Nguyen VVT, Gkouzioti V, Maass C, Verhaar MC, Vernooij RWM, van Balkom BWM. A systematic review of kidney-on-a-chip-based models to study human renal (patho-)physiology. Dis Model Mech. 2023;16(6). doi:10.1242/dmm.050113

25. Englezakis A, Gozalpour E, Kamran M, Fenner K, Mele E, Coopman K. Development of a hollow fibre-based renal module for active transport studies. J Artif Organs. 2021;24(4):473–484. doi:10.1007/S10047-021-01260-W

26. Jansen J, De Napoli IE, Fedecostante M, et al. Human proximal tubule epithelial cells cultured on hollow fibers: living membranes that actively transport organic cations. Scientific Reports 2015 5:1. 2015;5(1):1–12. doi:10.1038/srep16702

27. Homan KA, Kolesky DB, Skylar-Scott MA, et al. Bioprinting of 3D Convoluted Renal Proximal Tubules on Perfusable Chips. Scientific Reports 2016 6:1. 2016;6(1):1–13. doi:10.1038/srep34845

28. Lin NYC, Homan KA, Robinson SS, et al. Renal reabsorption in 3D vascularized proximal tubule models. Proc Natl Acad Sci U S A. 2019;116(12):5399–5404. doi:10.1073/pnas.1815208116

29. Vormann MK, Gijzen L, Hutter S, et al. Nephrotoxicity and Kidney Transport Assessment on 3D Perfused Proximal Tubules. AAPS Journal. 2018;20(5):1–11. doi:10.1208/S12248-018-0248-Z/FIGURES/5

30. Vormann MK, Tool LM, Ohbuchi M, et al. Modelling and Prevention of Acute Kidney Injury through Ischemia and Reperfusion in a Combined Human Renal Proximal Tubule/Blood Vessel-on-a-Chip. Kidney360. 2022;3(2):217–231. doi:10.34067/KID.0003622021/-/DCSUPPLEMENTAL

31. Gijzen L, Bokkers M, Hanamsagar R, et al. An immunocompetent human kidney on-a-chip model to study renal inflammation and immune-mediated injury. Biofabrication. 2024;17(1):015040. doi:10.1088/1758-5090/AD9FDF

32. Marschner JA, Martin L, Wilken G, Melica ME, Anders HJ. A “Kidney-on-the-Chip” Model Composed of Primary Human Tubular, Endothelial, and White Blood Cells. Methods in Molecular Biology. 2023;2664:107–121. doi:10.1007/978-1-0716-3179-9_8

33. Liu L, Zhang J, Bates S, et al. A methylation profile of in vitro immortalized human cell lines. Int J Oncol. 2005;26(1):275–285. doi:10.3892/ijo.26.1.275

34. Bens M, Vandewalle A. Cell models for studying renal physiology. Pflugers Arch. 2008;457(1):1–15. doi:10.1007/s00424-008-0507-4

35. Shi M, McCracken KW, Patel AB, et al. Human ureteric bud organoids recapitulate branching morphogenesis and differentiate into functional collecting duct cell types. Nature Biotechnology 2022 41:2. 2022;41(2):252–261. doi:10.1038/s41587-022-01429-5

36. Montalbetti N, Przepiorski AJ, Shi S, et al. Functional characterization of ion channels expressed in kidney organoids derived from human induced pluripotent stem cells. Am J Physiol Renal Physiol. 2022;323(4):F479–F491. doi:10.1152/AJPRENAL.00365.2021/ASSET/IMAGES/MEDIUM/F-00365-2021R01.PNG

37. Castrop H, Schießl IM. Physiology and pathophysiology of the renal Na-K-2Cl cotransporter (NKCC2). Am J Physiol Renal Physiol. 2014;307(9):F991–F1002. doi:10.1152/AJPRENAL.00432.2014/ASSET/IMAGES/LARGE/ZH20211474300001.JPEG

38. Markadieu N, Delpire E. Physiology and Pathophysiology of SLC12A1/2 transporters. Pflugers Arch. 2013;466(1):10.1007/s00424-013-1370-1375. doi:10.1007/S00424-013-1370-5

39. Butterworth MB. Regulation of the epithelial sodium channel (ENaC) by membrane trafficking. Biochim Biophys Acta. 2010;1802(12):1166. doi:10.1016/J.BBADIS.2010.03.010

40. Lever JE. REGULATION OF DOME FORMATION IN KIDNEY EPITHELIAL CELL CULTURES. Ann N Y Acad Sci. 1981;372(1). doi:10.1111/j.1749-6632.1981.tb15489.x

41. Latorre E, Kale S, Casares L, et al. Active superelasticity in three-dimensional epithelia of controlled shape. Nature. 2018;563(7730). doi:10.1038/s41586-018-0671-4

42. Biochemistr and, Cattaneo I, Condorelli L, et al. Cellular Physiology Cellular Physiology Cellular Physiology Cellular Physiology Cellular Physiology Shear Stress Reverses Dome Formation in Confluent Renal Tubular Cells. Original Paper Cell Physiol Biochem. 2011;28.

43. Oberleithner H, Vogel U, Kersting U. Madin-Darby canine kidney cells - I. Aldosterone-induced domes and their evaluation as a model system. Pflugers Arch. 1990;416(5). doi:10.1007/BF00382685

44. Rein JL, Heja S, Flores D, et al. Effect of luminal flow on doming of mpkCCD cells in a 3D perfusable kidney cortical collecting duct model. Am J Physiol Cell Physiol. 2020;318(1). doi:10.1152/ajpcell.00405.2019

45. Trietsch SJ, Naumovska E, Kurek D, et al. Membrane-free culture and real-time barrier integrity assessment of perfused intestinal epithelium tubes. Nature Communications 2017 8:1. 2017;8(1):1–8. doi:10.1038/s41467-017-00259-3

46. Mori H, Kuroda Y, Imamura S, et al. Hyponatremia and/or hyperkalemia in patients treated with the standard dose of trimethoprim-sulfamethoxazole. Intern Med. 2003;42(8):665–669. doi:10.2169/INTERNALMEDICINE.42.665

47. Lei H, Liu X, Zeng J, et al. Analysis of the Clinical Characteristics of Hyponatremia Induced by Trimethoprim/Sulfamethoxazole. Pharmacology. 2022;107(7-8):351–358. doi:10.1159/000523824

48. Tsapepas D, Chiles M, Babayev R, et al. Incidence of Hyponatremia with High-Dose Trimethoprim-Sulfamethoxazole Exposure. American Journal of Medicine. 2016;129(12):1322–1328. doi:10.1016/j.amjmed.2016.07.012

49. Marak C, Nunley M, Guddati AK, Kaushik P, Bannon M, Ashraf A. Severe hyponatremia due to trimethoprim-sulfamethoxazole-induced SIADH. SAGE Open Med Case Rep. 2022;10:2050313X221132654. doi:10.1177/2050313X221132654

50. Babayev R, Terner S, Chandra S, Radhakrishnan J, Mohan S. Trimethoprim-associated hyponatremia. Am J Kidney Dis. 2013;62(6):1188–1192. doi:10.1053/J.AJKD.2013.06.007

51. Muto S, Tsuruoka S, Miyata Y, Fujimura A, Kusano E. Effect of trimethoprim-sulfamethoxazole on Na and K+ transport properties in the rabbit cortical collecting duct perfused in vitro. Nephron Physiol. 2006;102(3-4). doi:10.1159/000089682

52. Hoorn EJ, Walsh SB, McCormick JA, et al. The calcineurin inhibitor tacrolimus activates the renal sodium chloride cotransporter to cause hypertension. Nat Med. 2011;17(10):1304–1309. doi:10.1038/NM.2497

53. Dimke H, Sparks MA, Thomson BR, Frische S, Coffman TM, Quaggin SE. Tubulovascular cross-talk by vascular endothelial growth factor a maintains peritubular microvasculature in kidney. Journal of the American Society of Nephrology. 2015;26(5). doi:10.1681/ASN.2014010060

54. Perretta-Tejedor N, Jafree DJ, Long DA. Endothelial-epithelial communication in polycystic kidney disease: Role of vascular endothelial growth factor signalling. Cell Signal. 2020;72. doi:10.1016/j.cellsig.2020.109624

55. Chen SJ, Lv LL, Liu BC, Tang RN. Crosstalk between tubular epithelial cells and glomerular endothelial cells in diabetic kidney disease. Cell Prolif. 2020;53(3). doi:10.1111/cpr.12763

